# Community context influences the conjugation efficiency of *E. coli*

**DOI:** 10.1101/2024.01.30.577951

**Authors:** Misshelle Bustamante, Floor Koopman, Jesper Martens, Jolanda K. Brons, Javier DelaFuente, Oscar P. Kuipers, Sander van Doorn, Marjon G.J. de Vos

## Abstract

In urinary tract infections different bacteria can live in a polymicrobial community, it is unknown how such community members affect the conjugation rate of uropathogenic *Escherichia coli*. We investigated the influence of the polymicrobial urinary tract infection (UTI) community context on the conjugation rate of *E. coli* isolates in artificial urine medium. Pairwise conjugation rate experiments were conducted between a donor *E. coli* strain containing pOXA-48 and six uropathogenic *E. coli* isolates in the presence and absence of five community members to elucidate their effect on the rate of conjugation. We found that the basal conjugation rates in the absence of community members are genotype dependent. Interestingly, bacterial interactions have an overall positive effect on *E. coli* conjugation rates. Particularly Gram-positive enterococcal species were found to enhance the conjugation rates of most uropathogenic *E. coli* isolates. We hypothesize that the nature and co-culture of the interactions is important for these increased conjugation rates in AUM.

## Introduction

Antimicrobial resistance (AMR) poses a significant challenge to global public health (Murray et al., 2022). The intense use of antibiotics has led to the emergence and spread of multidrug resistance in pathogenic bacteria (Polianciuc et al., 2020). This threatens the effectiveness of antibiotics, and therefore our ability to cure infections (Aslam et al., 2018; Prestinaci et al., 2015).

Bacteria can acquire antimicrobial resistance by horizontal exchange of genetic material among related or unrelated bacterial species, in a process referred to as ‘horizontal gene transfer’ or HGT (Hall et al., 2017; Ochman et al., 2000). The exchange of genetic material between microbes can occur in various ways, often by a process called conjugation (Furuya & Lowy, 2006). Conjugation involves the physical contact between donor and recipient cells and typically a self-transmissible or mobilizable plasmid (Ochman et al., 2000).

Conjugative or mobilizable plasmids are the most common transmission vectors for AMR genes (Ares-Arroyo et al., 2022; Boerlin & Reid-Smith, 2008; Partridge et al., 2018) and the major drivers of HGT within bacterial communities (Bottery, 2022). AMR genes and HGT have been observed within the human microbiome. A major hotspot for antibiotic resistance is, for instance, the gut microbiome of humans and animals (San Millan, 2018), where rich dynamics of plasmid transfer have been observed (Frazão et al., 2023).

Although the ecology (Smillie et al., 2011) and functioning of microbial communities are typically studied in one specific environment at a time, it is known that the rate of HGT is strongly dependent on the environment (Sessitsch et al., 2023). For instance, resource availability or other abiotic factors can significantly affect the rate at which HGT via conjugation occurs (Pallares-Vega et al., 2021). Biotic factors, such as the presence of ecological interaction partners, can affect the spread of conjugative plasmids within and between host species (Bottery, 2022), as well as the horizontal transmission of plasmids can be limited by bacterial diversity (Kottara et al., 2021). The microbial context can also determine the cost and benefits of conjugative plasmid maintenance (Sünderhauf et al., 2023). Moreover, ecological interactions can alter factors such as growth rate, and population densities, which together can affect the cost of plasmid carriage and conjugation rates (Duxbury et al., 2021).

It is an open question to what extent bacterial interactions affect the transfer of antibiotic resistance by HGT via conjugation in bacterial communities. Given that complex communities are difficult to study, we answer this question for a simple and tractable, yet relevant system of polymicrobial communities isolated from elderly patients diagnosed with urinary tract infections (UTIs) (Croxall et al., 2011). The prevalence of AMR in such communities is high (Croxall et al., 2011), and there has been an increase in AMR and multi-drug resistance in recent years (Trautner et al., 2022).

In such communities Gram-positive species live together with Gram-negative species (de Vos et al., 2017; Zandbergen et al., 2021), but the importance of Gram-positive species for UTIs is often overlooked. Yet, Gram-positive bacteria are an important cause of nosocomial infections (Cong et al., 2019; Furuno et al., 2005). Enterococci, for instance, have been shown to facilitate polymicrobial infections, leading to more complicated pathogenesis and poorer prognoses (Barshes et al., 2022; Chong et al., 2017), and they can compromise the efficacy of antimicrobial agents by promoting the colonization, proliferation, and persistence of diverse pathogenic bacteria (Xu et al., 2023). Furthermore, they can act as reservoirs for the transmission of antimicrobial resistance and virulence determinants (Coburn et al., 2007; Xu et al., 2021).

Here, we investigate the effect of ecological interactions between *E. coli* and other bacterial species isolated from polymicrobial UTI on the conjugation rate of pOXA-48 towards uropathogenic *E. coli*. pOXA-48 is a plasmid with a broad host range, carrying the resistance gene *bla*_*OXA-48*_ that confers resistance to multiple β-lactam antibiotics (Poirel et al., 2004, 2012), including carbapenems, which are last resort antibiotics used to treat multidrug resistant infections (Bradley et al., 1999; Papp-Wallace et al., 2011). It is an important conjugative plasmid in the clinical setting, known for its rapid dissemination within hospital patients and has a world-wide distribution (León-Sampedro et al., 2021; Pitout et al., 2019).

Specifically, we study the effect of ecological interactions on the conjugation rate in uropathogenic *E. coli* isolates, by performing pairwise conjugation assays in the presence of *Enterococcus faecium, Enterococcus faecalis, Staphylococcus simulans, Pseudomonas aeruginosa*, and *Proteus mirabilis* in artificial urine medium. All isolates were collected from elderly patients that were diagnosed with polymicrobial UTIs (Croxall et al., 2011).

## Materials and methods

### Bacterial isolates

Nine *E. coli* isolates were selected from a previous study where samples were collected from elderly patients diagnosed with polymicrobial urinary tract infections UTI (Croxall et al., 2011). They were selected based on their sensitivity to ampicillin. Initial conjugation experiments aimed at testing their ability to take up pOXA-48 plasmid resulted in six final uropathogenic *E. coli* isolates that were used in pairwise mating assays in the presence of UTI community members. Plasmid transfer in these isolates was confirmed by PCR with specific primers for pOXA-48 plasmid (see section DNA extraction and PCR in Materials and Methods).

The donor strain β3914 was a diaminopimelic acid (DAP) auxotrophic *E. coli* strain, exhibiting resistance to various antibiotics, including kanamycin (Roux et al., 2007) and harboring the pOXA-48 plasmid (Alonso-del Valle et al., 2021). This plasmid codes for the *bla*_*OXA-48*_ gene which confers resistance to β-lactam antibiotics (Poirel et al., 2004), including penicillins and carbapenems (Poirel et al., 2012).

The community members were collected from the same study as the uropathogenic *E. coli* isolates (Croxall et al., 2011), and were also selected upon their sensitivity to ampicillin. These belonged to three Gram-positive species: *E. faecium, E. faecalis, S. simulans*, and two Gram-negative species: *P. aeruginosa*, and *P. mirabilis*.

### Artificial urine medium (AUM)

We use a modified version of AUM (Brooks & Keevil, 1997; de Vos et al., 2017). It contained bacto peptone L37 1 g/L (Sigma), sodium bicarbonate 2.1 g/L (Roth), urea 7.5 g/L (Roth), sodium chloride 5.2 g/L (Sigma), sodium sulfate anhydrous 1.2 g/L, ammonium chloride 1.3 g/L (Sigma), and potassium dihydrogen phosphate 0.95 g/L added as solids; yeast extract 0.1 mL/L from 5g/100 mL stock, lactic acid 0.1 mL/L (Roth), citric acid 0.8 mL/L from 10g/20 mL stock, uric acid 7 mL/L from 1g/100mL in 1M NaOH stock, creatinine 16 mL/L from 5g/100mL stock, calcium chloride dihydrate 29.60 μL/L from 1g/10mL stock, iron(II) sulfate heptahydrate 12 μL/L from 10g/100mL stock, magnesium sulfate heptahydrate 2.45 mL/L from 1g/10mL stock were added as liquids.

### Conjugation on LB

This protocol was adapted from (Alonso-del Valle et al., 2021, 2023). Donor β3914 and recipient *E. coli* strains were streaked on CHROMagar plates supplemented with kanamycin 30 μg/mL (Sigma) and 300 μM DAP (Sigma), for the donor; and no antibiotic for the recipients, given that they were sensitive to most antibiotics used for the treatment of UTIs. The plates were incubated overnight at 37°C. The next day, 3 independent colonies were picked from each genotype and grown overnight in glass tubes with 2 mL of LB at 37°C and shaking at 200 rpm. Donor tubes were added 1 μL/mL of DAP.

The following day, 50 μL of the donors and 10 μL (5:1 donor to recipient ratio) were gently mixed in 0.5 mL tubes. The controls were isolated cultures of donor and recipients. The full 60 μL droplets were plated on independent LB agar (Sigma) plates without antibiotics but with 300 μM DAP (Sigma), and left to air-dry in the flow cabin, after which they were incubated at 37°C for 4 hours to recover transconjugants.

After incubation, a loop was used to scoop the biomass of the droplet, which was washed and resuspended in an Eppendorf tube with 2 mL of sterile 0.9% NaCl solution. The mix and controls were further diluted in serial dilutions from 10^1^ until 10^7^ using a 96-well plate: 200 μL of resuspension was added to the first well and the remaining wells contained 180 μL of 0.9% NaCl solution. Then, 20 μL from the first well was taken and mixed with the next well. This step was repeated for all columns of the well plate. A multichannel pipette was used to plate 10 μL from each dilution at the first quarter of an LB agar (Sigma) plate. The plate was tilted 90° to let the 8 droplets slide down until the end of the plate. Every donor and recipient mix, as well as the controls, were plated on LB plates with ampicillin 100 μg/mL and on LB plates with kanamycin 30 μg/mL (Sigma) without DAP, to make sure that neither donor nor recipients in isolation would grow.

After overnight incubation, glycerol stocks were made from the transconjugants. Given that this was a relatively crude, qualitative method to assess conjugation, a more quantifiable method was later applied to determine *E. coli* conjugation rates in the presence and absence of UTI community members.

### DNA extraction and PCR

To confirm plasmid transfer of the six *E. coli* isolates, DNA was extracted using the fast DNA extraction method described by (Brons et al., 2020). Primers for amplifying the resistance gene *bla*_*OXA-48*_ for β-lactam antibiotics on the pOXA-48 plasmid were adopted from (Poirel et al., 2004). A 20-mer forward primer, designated Oxa-48 Fw (5’-TTG GTG GCA TCG ATT ATC GG-3’) was combined with a 21-mer reverse primer, designated Oxa-48 Rev (5’-GAG CAC TTC TTT TGT GAT GGC-3’). This primer combination was tested and optimized.

PCR mixtures were prepared with the following components: 5.0 μl of 10x Roche buffer (Roche, Basel, Switzerland), 0.8 μl of 50mM MgCl_2_ (Merck, Darmstadt, Germany), 1.0 μl of 100% dimethyl sulfoxide (DMSO), 0.5 μl of 20 mg/ml bovine serum albumin (Merck, Darmstadt, Germany), 1.0 μl of 10 mM deoxyribonucleoside triphosphate mix, 1.0 μl of 10μM of each primer, and 0.2 μl of 5U/μl Taq DNA Polymerase (Roche, Basel, Switzerland). Molecular biology-grade water (Thermo Fisher Scientific, Waltham, United States) was added to a total volume of 50 μl in a 0.2-ml microfuge tube. Finally, 1.0 μl of template DNA was added. The mixtures were incubated in a Mastercycler Nexus PCR thermal cycler (Eppendorf, Hamburg, Germany) with the following program: initial denaturation of double-stranded DNA for 5 min at 95 °C; 35 cycles consisting of 1 min at 95 °C, 30 s. at 56 °C, and 2 min at 72 °C; and extension for 7 min at 72 °C.

All amplification products were analyzed by electrophoresis in 1.0% (wt/vol) agarose gels, followed by ethidium bromide staining (1.2 mg/ l ethidium bromide in 1× Tris-acetate-EDTA) (Mullis, 1990; Sambrook et al., 1989), destaining (1× Tris-acetate), and visualization under UV. Amplicons of 743 bp in size were detected, and no side products were observed, confirming plasmid transfer of the six *E. coli* isolates.

### Conjugation rate *E. coli* with community members on AUM

We performed a quantifiable method based on (Alonso-del Valle et al., 2023; DelaFuente et al., 2022; León-Sampedro et al., 2021) to determine the rates of conjugation of six *E. coli* isolates in the presence and absence of five polymicrobial UTI community members in AUM media. A scoop from -80°C glycerol stocks was taken to grow overnight cultures of donor, recipient *E. coli* and community members into glass tubes with 2 mL of 1x AUM. The donor strain β3914 with pOXA-48 plasmid was grown with 30 μg/mL kanamycin (Sigma) and 300 μM DAP (Sigma). The recipient and community member strain cultures had no additives. They were incubated for 24 hours at 37°C shaking at 200 rpm. For each experiment, the basal conjugation rate of a particular uropathogenic *E. coli* (donor and recipient only) was measured, as well as the community conjugation rate (donor, recipient and community member).

After 24 hours, the population size, as inferred from optical density measurements at 600 nm (OD600), was assessed for all strains and the cultures were diluted below 0.4 OD to obtain an accurate OD600 reading. They were then further diluted to obtain a 5:1:1 OD600 ratio between donor, recipient and community members with the OD600 values being 1, 0.2 and 0.2, respectively; except in the case of *E. faecium* and *E. faecalis*, where the OD was always lower than 0.2.

For the assessment of the basal conjugation rate of the *E. coli* in the absence of community members, 50 μL of donor and 50 μL of recipient were added and gently mixed in a 0.5 mL tube, keeping a 5:1 OD600 ratio. For the assessment of the community conjugation rate, 50 μL of donor and recipient were also added to a 0.5 mL tube with an additional volume from the community member culture that was dependent on the OD, preserving the 5:1:1 OD600 ratio of donor, recipient and community members. If the OD of the undiluted culture was <0.2, which was always the case with *E. faecium* and *E. faecalis*, then 50 μL of it was added to the tube. For mixing, vortex was avoided, and the tubes were gently struck several times. All 100 μL droplets were plated on 1x AUM agar plates containing DAP. These plates were prepared using 50% Micro agar (15 g/L) (Duchefa Biochemie), 50% 2x AUM and 300 μM DAP. The droplets were left to dry and incubated for conjugation at 37°C for 1 hour. This was performed in triplicates for every combination of donor + recipient + community member.

After 1 hour of incubation, the plates were removed from the incubator. A sterile toothpick was used to cut out the piece of agar with the droplet. Subsequently, the agar segment was crushed and resuspended in 1 mL sterile 0.9% NaCl solution. Each tube was inverted and gently shaken 30 times to wash off the cells from the agar. Further tenfold dilutions until 10^4^ were made before plating 100 μL of the re-suspensions to obtain countable colonies. Transconjugant colonies were obtained either at undiluted or 10^1^ diluted re-suspensions. The selective plates that were used to obtain donor, recipient and community members CFU/mL counts were LB agar (Sigma) with 30 μg/mL kanamycin and DAP, CHROMagar (without DAP) (Condalab), and CHROMagar (Condalab) with 100 μg/mL ampicillin; respectively. They were left overnight at 37°C and colonies were counted the next day.

The *E. coli* conjugation rate when community members are present was estimated using the formula: *T /* (*D* · *R* · Δ*t*) (Huisman et al., 2022; Lopatkin et al., 2016) where the CFU/mL of the transconjugants is represented by *T*; *D* are CFU/mL the donor, and *R* are CFU/mL of the recipient. The time in which conjugation took place is represented by *Δt*, and it was always 1 hour; the approximate time needed for pOXA-48 to produce transconjugants (León-Sampedro et al., 2021), yet keeping on-plate growth to a minimum.

### Spent media

Spent media were recovered from two UTI isolates; *E. faecalis* and *E. faecium* by inoculating bacterial glycerol stocks in 200 mL 1x AUM in Erlenmeyer flasks shaking at 200 rpm at 37°C for 48 hours. Afterwards, cultures were distributed into 50 mL culture tubes and centrifuged for 15 minutes at 4,800 g at room temperature. The resulting supernatants were filtered twice with bottle filter tops; 0.45 μm and 0.2 μm filters, respectively. To make sure that all bacteria were filtered out, spent medium was plated on CHROMagar plates, and incubated at 37°C for 24 hours, whereafter the plates showed no bacterial growth.

### Conjugation plates for spent media experiment

The spent media plates were prepared using 50% Micro agar (15 g/L) (Duchefa Biochemie), 25% spent media, and 25% 3x AUM (including 1x concentration AUM salts). Control plates consisted of 50% Micro agar (15 g/L), 25% 1x AUM, and 25% 3x AUM (including 1x concentration AUM salts). The end concentration of AUM in the spent media plates depended on how much nutrients were depleted by the bacteria, between 0,75x AUM (if all nutrients were consumed) and 1x AUM depending (if no nutrients were consumed). All plates contained 300 μM DAP to ensure survival of donor bacteria.

### Statistical analysis

All statistical analyses were performed in Rstudio2023.12.0+369 (2023.12.0+369). ANOVA was used on log transformed data to obtain the significant differences among the *E. coli* isolates, followed up by a Tukey-Kramer test. Pearson correlation coefficient measured the correlation between the basal conjugation rates for the six uropathogenic *E. coli* isolates and the effect of the community on the conjugation rate.

## Results

To investigate the impact of other community members isolated from polymicrobial UTIs on the conjugation rate in uropathogenic *E. coli*, we compared the pOXA-48 reception rate through conjugation of isolated uropathogenic *E. coli* with that of uropathogenic *E. coli* in the presence of other species isolated from polymicrobial UTIs.

Specifically, we performed conjugation experiments between donor strain β3914 and nine uropathogenic *E. coli* isolates, using plasmid pOXA-48; initially on LB media (Materials and Methods). Three out of these nine *E. coli* isolates didn’t take up the plasmid (Supplementary Table 1). Of the six uropathogenic *E. coli* isolates that did take the plasmid up (Figure 1 A-F, Materials and Methods), they did so with different conjugation efficiencies; indicated by the qualitative assessment of the maximum dilution that obtained transconjugants.

**Figure 1.**
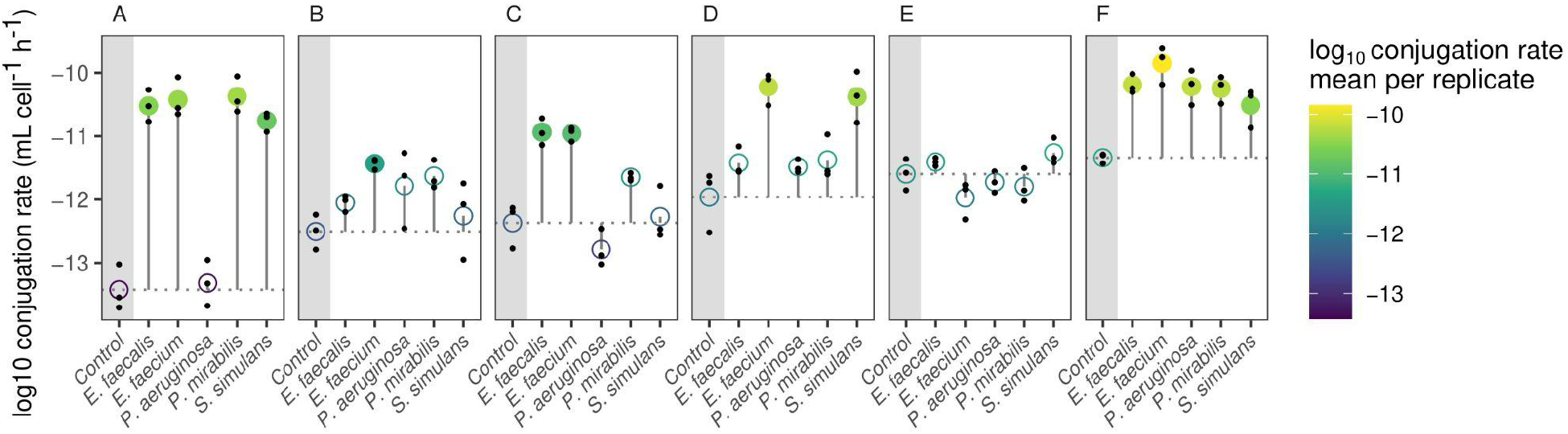
Conjugation rate of six *E. coli* isolates (A-F) in the presence and absence of other polymicrobial UTI isolates: *E. faecium, E. faecalis, S. simulans, P. aeruginosa* and *P. mirabilis*. Circles illustrate the mean per experiment. Controls represent the basal conjugation rate in the absence of community members. Solid circles indicate a significant difference between the controls and the experiments with each of the UTI community members (ANOVA on log-transformed data followed by Tukey-Kramer test).

With this in mind, we tested the differential effects on the conjugation rate of six *E. coli* isolates with donor strain β3914 in the absence and in the presence of five other UTI community members individually in AUM: *E. faecium, E. faecalis, S. simulans, P. aeruginosa* and *P. mirabilis*. Conjugation rate experiments were performed by bringing bacteria together in a droplet of conjugation mix, as described by (Alonso-del Valle et al., 2023; DelaFuente et al., 2022; León-Sampedro et al., 2021) on artificial urine medium (AUM) agar plates (Materials and Methods). For the community conjugation rate experiments, the donor and recipient *E. coli* pairs were spotted on the AUM agar plates, in the presence of one of the bacterial community members.

We found that the basal conjugation rates, in the absence of community members, differ between the six *E. coli* isolates. The conjugation rates in isolation range within two orders of magnitude between the different isolates; from 4.7 x 10^−14^ for *E. coli* isolate A, to 4.6 x 10^−12^ for *E. coli* isolate F (Figure 1). Assessing the effect of the bacterial interactions on the conjugation rate, we did not find that any of the other tested isolates inhibit the growth of either donor or recipient (Supplementary Table 2).

Rather, we interestingly found that bacterial interactions generally have a positive effect on the conjugation rates. Particularly the Gram-positive species *E. faecium* and *E. faecalis*, but also *S. simulans* contribute to this effect. For five of the six *E. coli* isolates, at least one Enterococcus species has a significant positive effect on the conjugation rate, whereas *S. simulans* has a positive effect on the conjugation rate of three of the *E. coli* isolates. The Gram-negative species *P. aeruginosa* and *P. mirabilis* generally have a less prominent effect. *P. mirabilis* alters the conjugation rate of two of the six *E. coli* species, whereas *P. aeruginosa* only affects the conjugation rate of one *E. coli* isolate (Figure 1; Supplementary Table 2).

The extent to which pairwise interactions changed the conjugation rate varied substantially between isolates. For instance, for *E. coli* isolate A, three UTI community members increased the conjugation rate by three orders of magnitude. For isolate C, the presence of both *Enterococcus* species increased the conjugation rate by two orders of magnitude. Yet, for isolate E we could not detect any significant changes to the conjugation rate due to the influence of the other community members (Figure 1). The magnitude of the variability of the conjugation rates was constant throughout the tested conditions (Supplementary Figure 1).

To assess whether the pairwise-interaction-dependent conjugation rate was correlated with the basal conjugation rate in isolation, we correlated the average effect of all pairwise interactions of each isolate to the basal conjugation rate. We do not find a significant correlation (Supplementary Figure 2 and 3). This shows that, regardless of the basal conjugation rate, the conjugation efficiency depends on the specific isolate, and the interaction with specific community members.

To investigate whether the increased conjugation rates, particularly due to the presence of Enterococci, were due to their exudates (for instance metabolic products produced), we tested whether the presence of conditioned media of *E. faecium* and *E. faecalis* can recapitulate the findings. Conjugation experiments were performed with three *E. coli* isolates (isolate A, B, F) on conditioned medium agar plates (Materials and Methods), with the spent media from the two Enterococci. Pairwise interactions were assessed by normalizing the conjugation rates on the conditioned medium agar plates to the conjugation rate in the absence of conditioned medium (Materials and Methods).

We found that the conditioned media experiments cannot explain the marked increase of the conjugation rate in the presence of *E. faecium* and *E. faecalis* (Figure 2). This indicates that the production of primary or secondary metabolites present in this cell-free medium conditioned by *E. faecium* and *E. faecalis* is unlikely to be the leading cause of the increased conjugation rates observed in the co-culture experiments. This suggests that Gram-positive bacteria affect the conjugative transfer or AMR in uropathogenic *E. coli*, in a manner that is likely dependent on the physical interaction of *E. coli* and the Gram-positive species.

**Figure 2.**
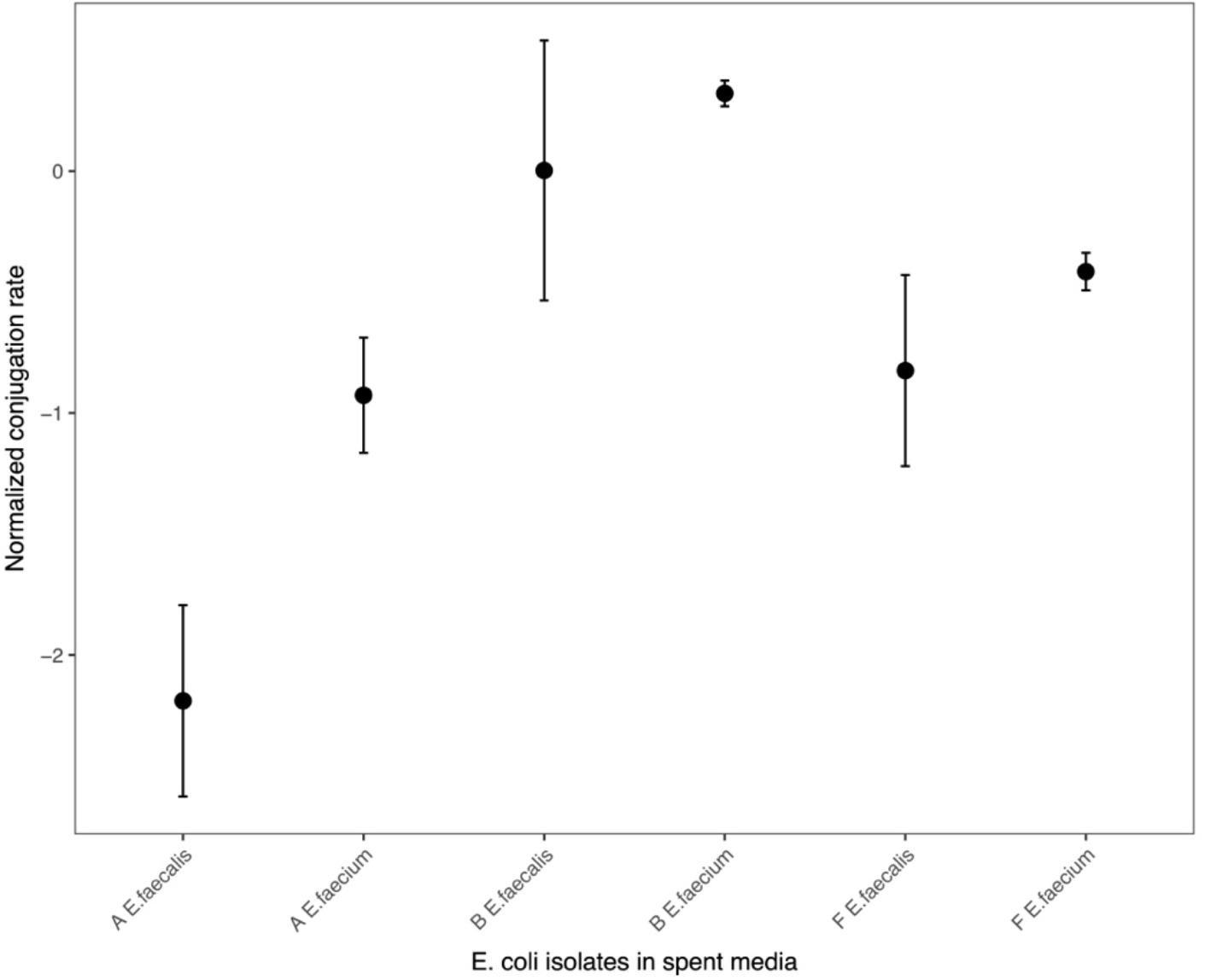
Conjugation rates of three of the *E. coli* isolates; A, B and F on spent media agar plates from *E. faecium* and *E. faecalis*. Black points represent the average conjugation rates, normalized to control (conjugation rate on agar plates with no spent media), and are given as log (fold change) from three replicates per experiment with standard deviation.

## Discussion

Nine uropathogenic *E. coli* isolates were initially selected for conjugation with pOXA-48. Only six of these *E. coli* isolates took up the plasmid, as verified by PCR with pOXA-48 specific primers (Materials and Methods). The fact that each of them had a unique basal conjugation rate indicates that there are host-dependent genetic background interactions that determine these rates, which is in accordance with other findings (Alonso-del Valle et al., 2023; Benz & Hall, 2023).

The conjugation rate experiments were performed in the low-nutrient artificial urine media (AUM) to recapitulate an environment that is closer to the *in vivo* environment of the uropathogens. Otherwise, similar conjugation experiments are often performed in Lysogeny broth LB (Alonso-del Valle et al., 2021; DelaFuente et al., 2022; León-Sampedro et al., 2021) or viande-levure VL (Duxbury et al., 2021; Huisman et al., 2022), which are rather rich media and yield mostly higher conjugation rates. The conjugation rates we find are similar to conjugation of pOXA-48 in one strain of *E. coli*, cultured under anaerobic conditions and in rather poor M9 minimal media (León-Sampedro et al., 2021). We therefore hypothesize that the similarly low basal conjugation rates result from the low nutrient environment. This hypothesis was confirmed by testing the ability to take-up pOXA-48 plasmid through conjugation via the rather crude qualitative droplet-droplet method in a rich LB medium (Materials and Methods; Supplementary Table 1).

Interestingly, the Gram-positive Enterococci hardly grow in the artificial urine medium (10^3^-10^4^ CFU/ml), these relatively low counts mimic their growth in urine (Flores-Mireles et al., 2015). Their cells are therefore present in low numbers in the conjugation rate experiments on the AUM agar plates. Yet, despite these low numbers they are able to induce this increase in conjugation rate.

Conditioned medium experiments indicate that metabolic compounds in the exudates of the co-cultured Enterococcal species are unlikely to be the cause of the increased transfer. Thus, population size effects mediated by such exudates of Enterococci are also unlikely to be involved (Keogh et al., 2016). Moreover, to limit such potential growth effects we performed the incubation step of the conjugation rate experiments for only one hour, where other studies often use longer incubation times (Alonso-del Valle et al., 2023). And lastly, because the conjugation efficiency is calculated based on the numbers of donors, recipients and transconjugants at the end of the one-hour conjugation incubation time (after the co-growth of the donor β3914, recipient *E. coli* and community member on the agar plate), we conclude that growth rate differences are not the cause of the marked increase in conjugation rate in the presence of these species.

Our findings suggest that the nature and co-culture of the interactions is important for these increased conjugation rates in artificial urine medium. This suggests that physical contact or proximity, potentially mediated by signaling molecules (Lin et al., 2021) between these species may play a role. Various types of cell-to-cell contact have shown to be involved in promoting the transfer of genetic material within species (Morawska & Kuipers, 2022). For example, pheromone-inducible aggregation substance of Enterococci, a virulence factor that promotes the aggregation and therefore proximity of Enterococci, has shown to have a positive effect on the conjugation efficiency of Enterococcus species (Waters & Dunny, 2001). Potentially such aggregation also increases the proximity of other conjugating species, such as *E. coli*, in mixed cultures.

One alternative hypothesis would be that the presence of some Gram-positive species is strengthening the interaction between the two *E. coli* strains (donor and recipient) as a sort of defense mechanism, limiting the direct interaction of the Gram-positive ‘intruder’ with the interacting *E. coli* strains. This is reminiscent of biofilm formation as a defense mechanism (Donlan & Costerton, 2002; Kumar et al., 2017).

The fact that the uropathogenic *E. coli* are conjugatable at these levels in the AUM medium suggests that the urinary tract and its urobiome, is a potential location where HGT takes place (Jones et al., 2021; Kuznetsova et al., 2022; Montelongo Hernandez et al., 2022; Wolfe & Brubaker, 2019).

Finally, our findings on the increased conjugation rates in uropathogenic *E. coli* in the presence of Gram-positive species underscore that ecological interactions are relevant for the conjugative transfer of AMR, also in a urine-like environment.

## Supplementary data

Supplementary data files are available as separate files.

## Acknowledgements

We kindly thank Thomas Hackl for the help with Figures 1 and Figure S1, and Alvaro San Millan for donating donor strain, plasmid and protocol and discussions, and for reading the manuscript.

## References

Alonso-del Valle, A., León-Sampedro, R., Rodríguez-Beltrán, J., DelaFuente, J., Hernández-García, M., Ruiz-Garbajosa, P., Cantón, R., Peña-Miller, R., & San Millán, A. (2021). Variability of plasmid fitness effects contributes to plasmid persistence in bacterial communities. Nature Communications, 12(1), 2653.

Alonso-del Valle, A., Toribio-Celestino, L., Quirant, A., Pi, C. T., DelaFuente, J., Canton, R., Rocha, E. P. C., Ubeda, C., Peña-Miller, R., & San Millan, A. (2023). Antimicrobial resistance level and conjugation permissiveness shape plasmid distribution in clinical enterobacteria. Proceedings of the National Academy of Sciences, 120(51).

Ares-Arroyo, M., Coluzzi, C., & Rocha, E. P. C. (2022). Origins of transfer establish networks of functional dependencies for plasmid transfer by conjugation. Nucleic Acids Research, 51(7), 3001–3016.

Aslam, B., Wang, W., Arshad, M. I., Khurshid, M., Muzammil, S., Rasool, M. H., Nisar, M. A., Alvi, R. F., Aslam, M. A., Qamar, M. U., Salamat, M. K. F., & Baloch, Z. (2018). Antibiotic resistance: A rundown of a global crisis. Infection and Drug Resistance, Volume 11, 1645–1658.

Barshes, N. R., Clark, N. J., Bidare, D., Dudenhoeffer, J. H., Mindru, C., & Rodriguez-Barradas, M. C. (2022). Polymicrobial foot infection patterns are common and associated with treatment failure. Open Forum Infectious Diseases, 9(10).

Benz, F., & Hall, A. R. (2023). Host-specific plasmid evolution explains the variable spread of clinical antibiotic-resistance plasmids. Proceedings of the National Academy of Sciences of the United States of America, 120(15), e2212147120.

Boerlin, P., & Reid-Smith, R. J. (2008). Antimicrobial resistance: Its emergence and transmission. Animal Health Research Reviews, 9(2), 115–126.

Bottery, M. J. (2022). Ecological dynamics of plasmid transfer and persistence in microbial communities. Current Opinion in Microbiology, 68, 102152.

Bradley, J. S., Garau, J., Lode, H., Rolston, K. V. I., Wilson, S. E., & Quinn, J. P. (1999). Carbapenems in clinical practice: A guide to their use in serious infection. International Journal of Antimicrobial Agents, 11(2), 93–100.

Brons, J. K., Vink, S. N., de Vos, M. G. J., Reuter, S., Dobrindt, U., & van Elsas, J. D. (2020). Fast identification of Escherichia coli in urinary tract infections using a virulence gene based PCR approach in a novel thermal cycler. Journal of Microbiological Methods, 169, 105799.

Chong, K. K. L., Tay, W. H., Janela, B., Yong, A. M. H., Liew, T. H., Madden, L., Keogh, D., Barkham, T. M. S., Ginhoux, F., Becker, D. L., & Kline, K. A. (2017). Enterococcus faecalis modulates immune activation and slows healing during wound infection. The Journal of Infectious Diseases, 216(12), 1644–1654.

Coburn, P. S., Baghdayan, A. S., Dolan, G., & Shankar, N. (2007). Horizontal transfer of virulence genes encoded on the Enterococcus faecalis pathogenicity island. Molecular Microbiology, 63(2), 530–544.

Cong, Y., Yang, S., & Rao, X. (2019). Vancomycin resistant Staphylococcus aureus infections: A review of case updating and clinical features. Journal of Advanced Research, 21, 169–176.

Croxall, G., Weston, V., Joseph, S., Manning, G., Cheetham, P., & McNally, A. (2011). Increased human pathogenic potential of Escherichia coli from polymicrobial urinary tract infections in comparison to isolates from monomicrobial culture samples. Journal of Medical Microbiology, 60(1), 102–109.

de Vos, M. G. J., Zagorski, M., McNally, A., & Bollenbach, T. (2017). Interaction networks, ecological stability, and collective antibiotic tolerance in polymicrobial infections. Proceedings of the National Academy of Sciences, 114(40), 10666–10671.

DelaFuente, J., Toribio-Celestino, L., Santos-Lopez, A., León-Sampedro, R., Valle, A. A., Costas, C., Hernández-García, M., Cui, L., Rodríguez-Beltrán, J., Bikard, D., Cantón, R., & Millan, A. S. (2022). Within-patient evolution of plasmid-mediated antimicrobial resistance. Nature Ecology & Evolution, 10.1038/s41559-022-01908–7.

Donlan, R. M., & Costerton, J. W. (2002). Biofilms: Survival mechanisms of clinically relevant microorganisms. Clinical Microbiology Reviews, 15(2), 167–193.

Duxbury, S. J. N., Alderliesten, J. B., Zwart, M. P., Stegeman, A., Fischer, E. A. J., & de Visser, J. A. G. M. (2021). Chicken gut microbiome members limit the spread of an antimicrobial resistance plasmid in Escherichia coli. Proceedings of the Royal Society B: Biological Sciences, 288(1962), 20212027.

Flores-Mireles, A. L., Walker, J. N., Caparon, M., & Hultgren, S. J. (2015). Urinary tract infections: Epidemiology, mechanisms of infection and treatment options. Nature Reviews. Microbiology, 13(5), 269–284.

Frazão, N., Seixas, E., Barreto, H. C., Mischler, M., Güleresi, D., & Gordo, I. (2023). Massive lateral gene transfer under strain coexistence in the gut [Preprint]. Evolutionary Biology.

Furuno, J. P., Perencevich, E. N., Johnson, J. A., Wright, M.-O., McGregor, J. C., Morris, J. G., Strauss, S. M., Roghman, M.-C., Nemoy, L. L., Standiford, H. C., Hebden, J. N., & Harris, A. D. (2005). Methicillin-resistant Staphylococcus aureus and vancomycin-resistant Enterococci co-colonization. Emerging Infectious Diseases, 11(10), 1539–1544.

Furuya, E. Y., & Lowy, F. D. (2006). Antimicrobial-resistant bacteria in the community setting. Nature Reviews Microbiology, 4(1), 36–45.

Hall, J. P. J., Brockhurst, M. A., & Harrison, E. (2017). Sampling the mobile gene pool: Innovation via horizontal gene transfer in bacteria. Philosophical Transactions of the Royal Society B: Biological Sciences, 372(1735), 20160424.

Huisman, J. S., Benz, F., Duxbury, S. J. N., de Visser, J. A. G. M., Hall, A. R., Fischer, E. A. J., & Bonhoeffer, S. (2022). Estimating plasmid conjugation rates: A new computational tool and a critical comparison of methods. Plasmid, 121, 102627.

Jones, J., Murphy, C. P., Sleator, R. D., & Culligan, E. P. (2021). The urobiome, urinary tract infections, and the need for alternative therapeutics. Microbial Pathogenesis, 161, 105295.

Keogh, D., Tay, W. H., Ho, Y. Y., Dale, J. L., Chen, S., Umashankar, S., Williams, R. B. H., Chen, S. L., Dunny, G. M., & Kline, K. A. (2016). Enterococcal metabolite cues facilitate interspecies niche modulation and polymicrobial infection. Cell Host & Microbe, 20(4), 493–503.

Kottara, A., Carrilero, L., Harrison, E., Hall, J. P. J., & Brockhurst, M. A. (2021). The dilution effect limits plasmid horizontal transmission in multispecies bacterial communities. Microbiology, 167(9).

Kumar, A., Alam, A., Rani, M., Ehtesham, N. Z., & Hasnain, S. E. (2017). Biofilms: Survival and defense strategy for pathogens. International Journal of Medical Microbiology, 307(8), 481–489.

Kuznetsova, M. V., Maslennikova, I. L., Pospelova, J. S., Žgur Bertok, D., & Starčič Erjavec, M. (2022). Differences in recipient ability of uropathogenic Escherichia coli strains in relation with their pathogenic potential. Infection, Genetics and Evolution, 97, 105160.

León-Sampedro, R., DelaFuente, J., Díaz-Agero, C., Crellen, T., Musicha, P., Rodríguez-Beltrán, J., de la Vega, C., Hernández-García, M., López-Fresneña, N., Ruiz-Garbajosa, P., Cantón, R., Cooper, B. S., & Millán, Á. S. (2021). Pervasive transmission of a carbapenem resistance plasmid in the gut microbiota of hospitalised patients. Nature Microbiology, 6(5), 606–616.

Lin, Y.-C., Chen, E. H.-L., Chen, R. P.-Y., Dunny, G. M., Hu, W.-S., & Lee, K.-T. (2021). Biofilm-related infections: Bridging the gap between clinical management and fundamental aspects of recalcitrance toward antibiotics. Applied and Environmental Microbiology, 87(13), e00442–21.

Lopatkin, A. J., Huang, S., Smith, R. P., Srimani, J. K., Sysoeva, T. A., Bewick, S., Karig, D., & You, L. (2016). Antibiotics as a selective driver for conjugation dynamics. Nature Microbiology, 1(6), 16044.

Montelongo Hernandez, C., Putonti, C., & Wolfe, A. J. (2022). Profiling the plasmid conjugation potential of urinary Escherichia coli. Microbial Genomics, 8(5), mgen000814.

Morawska, L. P., & Kuipers, O. P. (2022). Cell-to-cell non-conjugative plasmid transfer between Bacillus subtilis and lactic acid bacteria. Microbial Biotechnology, 16(4), 784–798.

Mullis, K. B. (1990). The unusual origin of the polymerase chain reaction. Scientific American, 262(4), 56–61, 64–65.

Ochman, H., Lawrence, J. G., & Groisman, E. A. (2000). Lateral gene transfer and the nature of bacterial innovation. Nature, 405, 299–304.

Pallares-Vega, R., Macedo, G., Brouwer, M. S. M., Hernandez Leal, L., van der Maas, P., van Loosdrecht, M. C. M., Weissbrodt, D. G., Heederik, D., Mevius, D., & Schmitt, H. (2021). Temperature and nutrient limitations decrease transfer of conjugative IncP-1 plasmid pKJK5 to wild Escherichia coli strains. Frontiers in Microbiology, 12.

Papp-Wallace, K. M., Endimiani, A., Taracila, M. A., & Bonomo, R. A. (2011). Carbapenems: Past, present, and future ▿. Antimicrobial Agents and Chemotherapy, 55(11), 4943–4960.

Partridge, S. R., Kwong, S. M., Firth, N., & Jensen, S. O. (2018). Mobile genetic elements associated with antimicrobial resistance. Clinical Microbiology Reviews, 31(4), e00088–17.

Pitout, J. D. D., Peirano, G., Kock, M. M., Strydom, K.-A., & Matsumura, Y. (2019). The global ascendency of OXA-48-type Carbapenemases. Clinical Microbiology Reviews, 33(1), e00102–19.

Poirel, L., Bonnin, R. A., & Nordmann, P. (2012). Genetic features of the widespread plasmid coding for the Carbapenemase OXA-48. Antimicrobial Agents and Chemotherapy, 56(1), 559–562.

Poirel, L., Héritier, C., Tolün, V., & Nordmann, P. (2004). Emergence of Oxacillinase-mediated resistance to imipenem in Klebsiella pneumoniae. Antimicrobial Agents and Chemotherapy, 48(1), 15–22.

Polianciuc, S. I., Gurzău, A. E., Kiss, B., Ştefan, M. G., & Loghin, F. (2020). Antibiotics in the environment: Causes and consequences. Medicine and Pharmacy Reports, 93(3), 231–240.

Prestinaci, F., Pezzotti, P., & Pantosti, A. (2015). Antimicrobial resistance: A global multifaceted phenomenon. Pathogens and Global Health, 109(7), 309–318.

Roux, F. L., Binesse, J., Saulnier, D., & Mazel, D. (2007). Construction of a <i>Vibrio splendidus<i> mutant lacking the metalloprotease gene <i>vsm<i> by use of a novel counterselectable suicide vector. Applied and Environmental Microbiology, 73(3), 777.

Sambrook, J., Fritsch, E. F., & Maniatis, T. (1989). Molecular Cloning: A Laboratory Manual (2nd ed.). Cold Spring Harbor Laboratory Press.

San Millan, A. (2018). Evolution of plasmid-mediated antibiotic resistance in the clinical context. Trends in Microbiology, 26(12), 978–985.

Sessitsch, A., Wakelin, S., Schloter, M., Maguin, E., Cernava, T., Champomier-Verges, M.-C., Charles, T. C., Cotter, P. D., Ferrocino, I., Kriaa, A., Lebre, P., Cowan, D., Lange, L., Kiran, S., Markiewicz, L., Meisner, A., Olivares, M., Sarand, I., Schelkle, B., … Kostic, T. (2023). Microbiome interconnectedness throughout environments with major consequences for healthy people and a healthy planet. Microbiology and Molecular Biology Reviews, 87(3), e00212–22.

Smillie, C. S., Smith, M. B., Friedman, J., Cordero, O. X., David, L. A., & Alm, E. J. (2011). Ecology drives a global network of gene exchange connecting the human microbiome. Nature, 480, Article 7376.

Sünderhauf, D., Klümper, U., Gaze, W. H., Westra, E. R., & van Houte, S. (2023). Interspecific competition can drive plasmid loss from a focal species in a microbial community. The ISME Journal, 1–9.

Trautner, B. W., Kaye, K. S., Gupta, V., Mulgirigama, A., Mitrani-Gold, F. S., Scangarella-Oman, N. E., Yu, K., Ye, G., & Joshi, A. V. (2022). Risk factors associated with antimicrobial resistance and adverse short-term health outcomes among adult and adolescent female outpatients with uncomplicated urinary tract infection. Open Forum Infectious Diseases, 9(12), ofac623.

Waters, C. M., & Dunny, G. M. (2001). Analysis of functional domains of the Enterococcus faecalis pheromone-induced surface protein aggregation substance. Journal of Bacteriology, 183(19), 5659–5667.

Wolfe, A. J., & Brubaker, L. (2019). Urobiome updates: Advances in urinary microbiome research. Nature Reviews. Urology, 16(2), 73–74.

Xu, W., Fang, Y., Hu, Q., & Zhu, K. (2021). Emerging risks in food: Probiotic Enterococci pose a threat to public health through the food chain. Foods, 10(11), 2846.

Xu, W., Fang, Y., & Zhu, K. (2023). Enterococci facilitate polymicrobial infections. Trends in Microbiology.

Zandbergen, L. E., Halverson, T., Brons, J. K., Wolfe, A. J., & de Vos, M. G. J. (2021). The good and the bad: Ecological interaction measurements between the urinary microbiota and uropathogens. Frontiers in Microbiology, 12, 659450.

