## Supplementary data for "Community context influences the conjugation efficiency of *E. coli*"

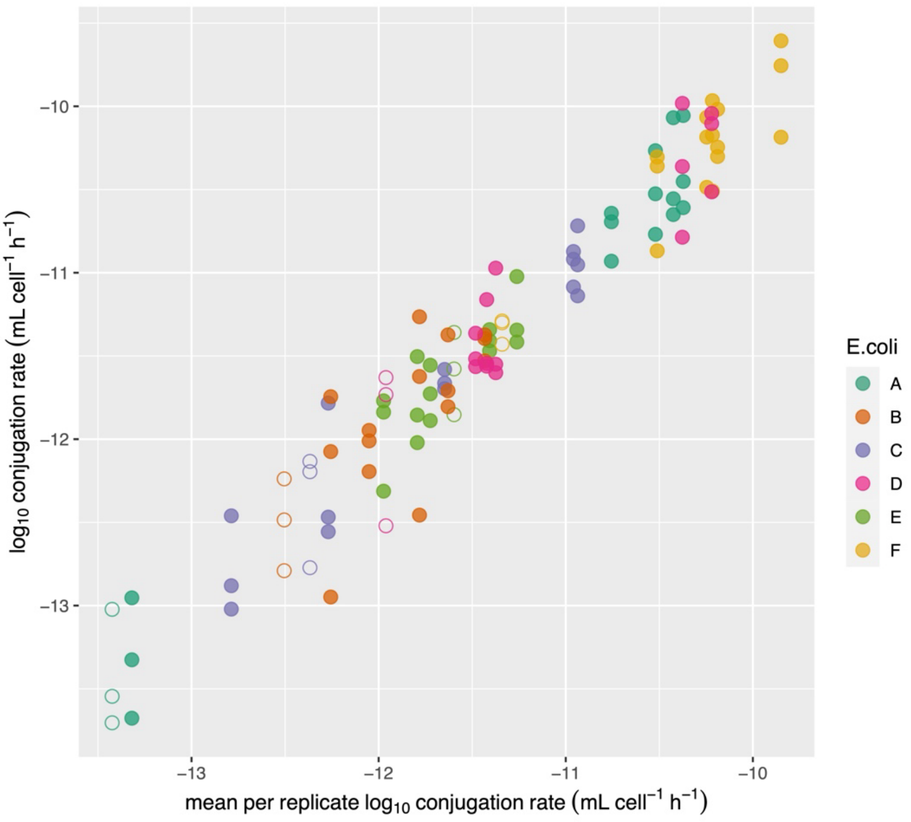


**Supplementary Figure 1.** Distribution of conjugation rate of pOXA-48 for six uropathogenic *E. coli* isolates (A-F) in the presence (solid circles) and absence (control; empty circles) of UTI community members: *E. faecium*, *E. faecalis*, *S. simulans, P. aeruginosa* and *P. mirabilis*. This graph shows that the variance of the conjugation rate in the different isolates, and in the presence and absence of community members, was rather constant.

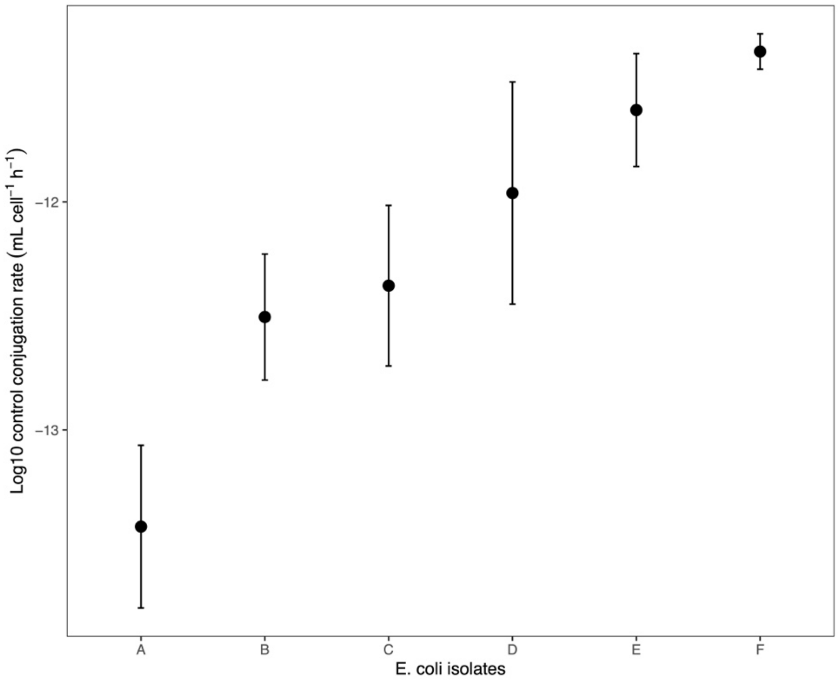


**Supplementary Figure 2.** Basal conjugation rates for six uropathogenic *E. coli* isolates (A-F) in the absence of the UTI community members. Circles represent the average conjugation rate of three replicates per isolate, error bars correspond to standard deviations.

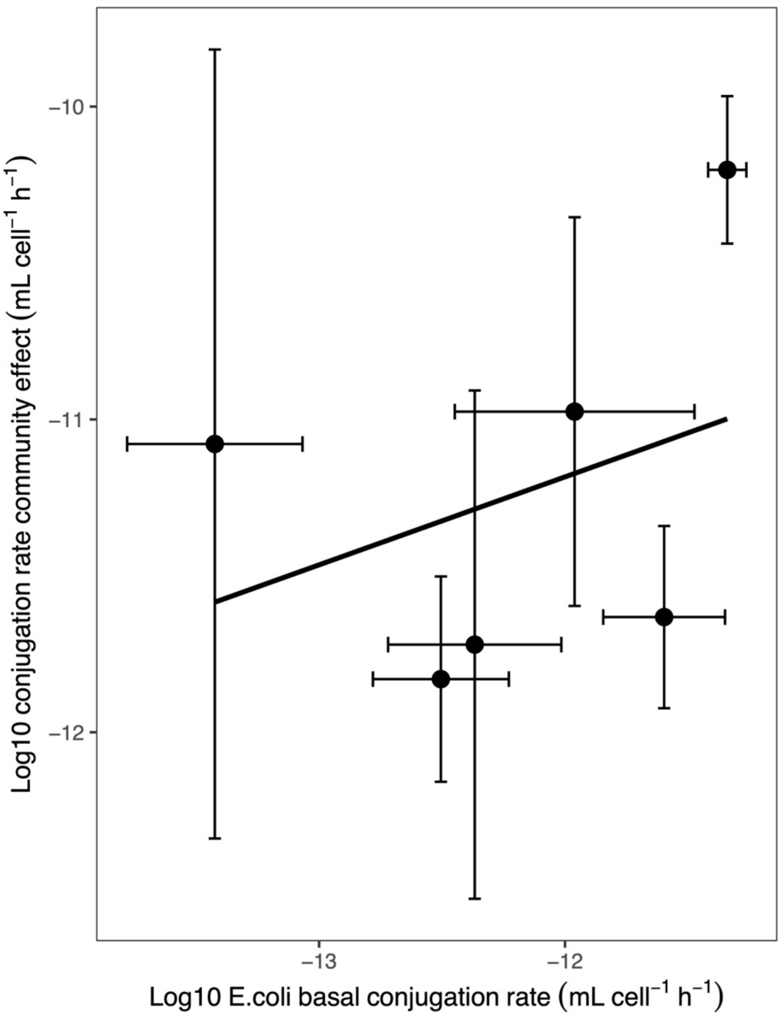


**Supplementary Figure 3**. Basal conjugation rates for six uropathogenic *E. coli* isolates against the conjugation rate effect of the community. Black circles represent the average basal conjugation rate of three replicates per *E. coli* isolate against the average of the conjugation rates of each UTI community member: *E. faecium*, *E. faecalis*, *S. simulans, P. aeruginosa* and *P. mirabilis*. Error bars correspond to standard deviation. The correlation of the basal conjugation rates, with the mean effect of all community members was not significant (pearson correlation coefficient 0.1154, p-value=0.5101).

**Supplementary Table 1**. Conjugation experiment on LB. Nine *E. coli* isolates were combined with donor strain β3914 containing pOXA-48 plasmid on LB agar, in triplicates. Dilutions from $10^1$ until $10^7$ were used to obtain transconjugants. This table shows that the isolates that took up the plasmid, did so with different conjugation efficiencies; indicated by the maximum dilution possible to obtain transconjugants on the selective plate. Interestingly, the rank of conjugation efficiency in LB and AUM is not equal.

| Strain_selected | Max_dilution |
|---|---|
| A | 1E+06 |
| A | 1E+06 |
| A | 1E+06 |
| B | 1E+02 |
| B | 1E+02 |
| B | 1E+02 |
| C | 1E+03 |
| C | 1E+03 |
| C | 1E+03 |
| D | 1E+05 |
| D | 1E+05 |
| D | 1E+05 |
| E | 1E+05 |
| E | 1E+05 |
| E | 1E+05 |
| F | 1E+07 |
| F | 1E+07 |
| F | 1E+07 |
| N/A | 0E+00 |
| N/A | 0E+00 |
| N/A | 0E+00 |
| N/A | 0E+00 |
| N/A | 0E+00 |
| N/A | 0E+00 |
| N/A | 0E+00 |
| N/A | 0E+00 |
| N/A | 0E+00 |

**Supplementary Table 2**. Individual CFU/mL counts of donors, recipients and transconjugants for the six uropathogenic *E. coli* with and without community member conjugation experiments on AUM agar plates. The triplicates of every experiment in the absence (control) and presence of every UTI community member: *E. faecium*, *E. faecalis*, *S. simulans, P. aeruginosa* and *P. mirabilis* are listed.

| Strain | Date | Experiment | Donor (CFU/mL) | Recipient CFU/mL | Transconjugant (CFU/mL) |
|---|---|---|---|---|---|
| A | 17_05_23 | Control | 1,09E+07 | 2,90E+07 | 3,00E+01 |
| A | 17_05_23 | Control | 2,25E+07 | 2,25E+07 | 1,00E+01 |
| A | 17_05_23 | Control | 2,65E+07 | 2,65E+07 | 2,00E+01 |
| A | 17_05_23 | *Enterococcus faecium* | 3,39E+07 | 3,80E+06 | 2,88E+03 |
| A | 17_05_23 | *Enterococcus faecium* | 2,94E+07 | 4,00E+06 | 3,27E+03 |
| A | 17_05_23 | *Enterococcus faecium* | 1,16E+07 | 3,00E+06 | 2,97E+03 |
| A | 17_05_23 | *Enterococcus faecalis* | 2,93E+07 | 6,60E+06 | 3,29E+03 |
| A | 17_05_23 | *Enterococcus faecalis* | 2,14E+07 | 8,00E+06 | 5,10E+03 |
| A | 17_05_23 | *Enterococcus faecalis* | 2,75E+07 | 3,90E+06 | 5,80E+03 |
| A | 17_05_23 | *Staphylococcus simulans* | 4,31E+07 | 3,10E+06 | 3,04E+03 |
| A | 17_05_23 | *Staphylococcus simulans* | 4,44E+07 | 4,60E+06 | 4,13E+03 |
| A | 17_05_23 | *Staphylococcus simulans* | 4,08E+07 | 5,10E+06 | 2,44E+03 |
| A | 17_05_23 | *Pseudomonas aeruginosa* | 2,38E+07 | 2,67E+07 | 3,00E+01 |
| A | 17_05_23 | *Pseudomonas aeruginosa* | 2,09E+07 | 2,58E+07 | 6,00E+01 |
| A | 17_05_23 | *Pseudomonas aeruginosa* | 2,21E+07 | 2,15E+07 | 1,00E+01 |
| A | 17_05_23 | *Proteus mirabilis* | 1,59E+07 | 6,20E+06 | 3,48E+03 |
| A | 17_05_23 | *Proteus mirabilis* | 1,20E+07 | 3,20E+06 | 3,38E+03 |
| A | 17_05_23 | *Proteus mirabilis* | 1,15E+07 | 3,50E+06 | 9,90E+02 |
| B | 26_10_23 | Control | 1,54E+07 | 4,50E+06 | 4,00E+01 |
| B | 26_10_23 | Control | 1,39E+07 | 4,40E+06 | 2,00E+01 |
| B | 26_10_23 | Control | 1,21E+07 | 5,10E+06 | 1,00E+01 |
| B | 26_10_23 | *Enterococcus faecium* | 1,38E+07 | 2,70E+06 | 1,10E+02 |
| B | 26_10_23 | *Enterococcus faecium* | 1,16E+07 | 1,50E+06 | 7,00E+01 |
| B | 26_10_23 | *Enterococcus faecium* | 1,26E+07 | 1,50E+06 | 8,00E+01 |
| B | 26_10_23 | *Enterococcus faecalis* | 1,99E+07 | 1,38E+07 | 3,10E+02 |
| B | 26_10_23 | *Enterococcus faecalis* | 1,59E+07 | 2,19E+07 | 3,40E+02 |
| B | 26_10_23 | *Enterococcus faecalis* | 1,27E+07 | 2,96E+07 | 2,40E+02 |
| B | 26_10_23 | *Staphylococcus simulans* | 7,50E+06 | 9,50E+06 | 6,00E+01 |
| B | 26_10_23 | *Staphylococcus simulans* | 1,96E+07 | 1,36E+07 | 3,00E+01 |
| B | 26_10_23 | *Staphylococcus simulans* | 8,00E+06 | 1,32E+07 | 1,90E+02 |
| B | 26_10_23 | *Pseudomonas aeruginosa* | 1,30E+07 | 8,80E+06 | 4,00E+01 |
| B | 26_10_23 | *Pseudomonas aeruginosa* | 5,30E+06 | 1,19E+07 | 1,50E+02 |
| B | 26_10_23 | *Pseudomonas aeruginosa* | 3,20E+06 | 4,60E+06 | 8,00E+01 |
| B | 26_10_23 | *Proteus mirabilis* | 9,60E+06 | 6,40E+06 | 1,20E+02 |
| B | 26_10_23 | *Proteus mirabilis* | 5,40E+06 | 7,00E+06 | 1,60E+02 |
| B | 26_10_23 | *Proteus mirabilis* | 9,30E+06 | 1,03E+07 | 1,50E+02 |
| C | 31_10_23 | Control | 5,10E+06 | 8,00E+06 | 3,00E+01 |
| C | 31_10_23 | Control | 8,80E+06 | 1,07E+07 | 6,00E+01 |
| C | 31_10_23 | Control | 8,00E+06 | 1,48E+07 | 2,00E+01 |
| C | 31_10_23 | *Enterococcus faecium* | 1,77E+07 | 8,60E+06 | 1,25E+03 |
| C | 31_10_23 | *Enterococcus faecium* | 1,32E+07 | 9,60E+06 | 1,70E+03 |
| C | 31_10_23 | *Enterococcus faecium* | 1,38E+07 | 1,13E+07 | 1,88E+03 |
| C | 31_10_23 | *Enterococcus faecalis* | 1,12E+07 | 1,00E+07 | 1,25E+03 |
| C | 31_10_23 | *Enterococcus faecalis* | 1,44E+07 | 1,30E+07 | 1,36E+03 |
| C | 31_10_23 | *Enterococcus faecalis* | 7,80E+06 | 7,10E+06 | 1,06E+03 |
| C | 31_10_23 | *Staphylococcus simulans* | 4,50E+06 | 8,00E+06 | 1,00E+01 |
| C | 31_10_23 | *Staphylococcus simulans* | 4,20E+06 | 7,00E+06 | 1,00E+01 |
| C | 31_10_23 | *Staphylococcus simulans* | 4,40E+06 | 6,90E+06 | 5,00E+01 |
| C | 31_10_23 | *Pseudomonas aeruginosa* | 5,30E+06 | 1,09E+07 | 2,00E+01 |
| C | 31_10_23 | *Pseudomonas aeruginosa* | 4,90E+06 | 1,55E+07 | 1,00E+01 |
| C | 31_10_23 | *Pseudomonas aeruginosa* | 7,50E+06 | 1,40E+07 | 1,00E+01 |
| C | 31_10_23 | *Proteus mirabilis* | 1,63E+07 | 9,90E+06 | 3,50E+02 |
| C | 31_10_23 | *Proteus mirabilis* | 1,82E+07 | 6,70E+06 | 3,20E+02 |
| C | 31_10_23 | *Proteus mirabilis* | 1,93E+07 | 9,80E+06 | 3,80E+02 |

| | | | | | |
|---|---|---|---|---|---|
| D | 29_06_23 | Control | 9,00E+06 | 1,80E+06 | 3,00E+01 |
| D | 29_06_23 | Control | 2,03E+07 | 4,90E+06 | 3,00E+01 |
| D | 29_06_23 | Control | 1,28E+07 | 4,00E+06 | 1,20E+02 |
| D | 29_06_23 | *Enterococcus faecium* | 1,72E+07 | 3,10E+06 | 4,20E+03 |
| D | 29_06_23 | *Enterococcus faecium* | 1,42E+07 | 3,50E+06 | 4,50E+03 |
| D | 29_06_23 | *Enterococcus faecium* | 1,66E+07 | 5,90E+06 | 3,00E+03 |
| D | 29_06_23 | *Enterococcus faecalis* | 3,47E+07 | 8,40E+06 | 8,00E+02 |
| D | 29_06_23 | *Enterococcus faecalis* | 2,26E+07 | 7,70E+06 | 1,20E+03 |
| D | 29_06_23 | *Enterococcus faecalis* | 3,77E+07 | 9,30E+06 | 1,00E+03 |
| D | 29_06_23 | *Staphylococcus simulans* | 1,41E+07 | 3,50E+06 | 2,00E+03 |
| D | 29_06_23 | *Staphylococcus simulans* | 7,80E+06 | 4,70E+06 | 3,50E+03 |
| D | 29_06_23 | *Staphylococcus simulans* | 1,01E+07 | 3,20E+06 | 1,00E+03 |
| D | 29_06_23 | *Pseudomonas aeruginosa* | 1,41E+07 | 3,50E+06 | 1,50E+02 |
| D | 29_06_23 | *Pseudomonas aeruginosa* | 7,80E+06 | 4,70E+06 | 1,00E+02 |
| D | 29_06_23 | *Pseudomonas aeruginosa* | 1,01E+07 | 3,20E+06 | 1,40E+02 |
| D | 29_06_23 | *Proteus mirabilis* | 1,34E+07 | 7,00E+06 | 1,00E+03 |
| D | 29_06_23 | *Proteus mirabilis* | 1,30E+07 | 9,20E+06 | 3,00E+02 |
| D | 29_06_23 | *Proteus mirabilis* | 1,94E+07 | 1,28E+07 | 7,00E+02 |
| E | 22_06_23 | Control | 6,40E+06 | 1,00E+07 | 2,80E+02 |
| E | 22_06_23 | Control | 6,10E+06 | 1,52E+07 | 1,30E+02 |
| E | 22_06_23 | Control | 1,40E+07 | 9,20E+06 | 3,40E+02 |
| E | 22_06_23 | *Enterococcus faecium* | 4,10E+07 | 2,01E+07 | 1,40E+03 |
| E | 22_06_23 | *Enterococcus faecium* | 3,63E+07 | 2,83E+07 | 5,00E+02 |
| E | 22_06_23 | *Enterococcus faecium* | 3,08E+07 | 3,80E+07 | 1,70E+03 |
| E | 22_06_23 | *Enterococcus faecalis* | 3,27E+07 | 3,61E+07 | 4,00E+03 |
| E | 22_06_23 | *Enterococcus faecalis* | 2,92E+07 | 3,86E+07 | 4,40E+03 |
| E | 22_06_23 | *Enterococcus faecalis* | 3,32E+07 | 4,04E+07 | 6,10E+03 |
| E | 22_06_23 | *Staphylococcus simulans* | 2,36E+07 | 1,57E+07 | 1,42E+03 |
| E | 22_06_23 | *Staphylococcus simulans* | 3,73E+07 | 6,70E+06 | 2,37E+03 |
| E | 22_06_23 | *Staphylococcus simulans* | 3,70E+07 | 9,50E+06 | 1,59E+03 |
| E | 22_06_23 | *Pseudomonas aeruginosa* | 2,08E+07 | 3,46E+07 | 9,30E+02 |
| E | 22_06_23 | *Pseudomonas aeruginosa* | 1,53E+07 | 2,89E+07 | 1,23E+03 |
| E | 22_06_23 | *Pseudomonas aeruginosa* | 1,76E+07 | 3,40E+07 | 1,12E+03 |
| E | 22_06_23 | *Proteus mirabilis* | 3,87E+07 | 2,44E+07 | 9,00E+02 |
| E | 22_06_23 | *Proteus mirabilis* | 2,59E+07 | 2,49E+07 | 9,00E+02 |
| E | 22_06_23 | *Proteus mirabilis* | 2,80E+07 | 1,71E+07 | 1,50E+03 |
| F | 12_05_23 | Control | 5,90E+06 | 2,38E+07 | 7,20E+02 |
| F | 12_05_23 | Control | 1,12E+07 | 2,45E+07 | 1,02E+03 |
| F | 12_05_23 | Control | 1,30E+07 | 1,43E+07 | 9,30E+02 |
| F | 12_05_23 | *Enterococcus faecium* | 6,70E+06 | 4,00E+06 | 4,70E+03 |
| F | 12_05_23 | *Enterococcus faecium* | 6,10E+06 | 5,90E+06 | 8,90E+03 |
| F | 12_05_23 | *Enterococcus faecium* | 1,07E+07 | 4,30E+06 | 3,00E+03 |
| F | 12_05_23 | *Enterococcus faecalis* | 3,35E+07 | 1,60E+07 | 5,14E+04 |
| F | 12_05_23 | *Enterococcus faecalis* | 3,83E+07 | 2,05E+07 | 4,46E+04 |
| F | 12_05_23 | *Enterococcus faecalis* | 4,02E+07 | 1,63E+07 | 3,27E+04 |
| F | 12_05_23 | *Staphylococcus simulans* | 4,31E+07 | 7,20E+06 | 4,20E+03 |
| F | 12_05_23 | *Staphylococcus simulans* | 3,70E+07 | 6,00E+06 | 9,70E+03 |
| F | 12_05_23 | *Staphylococcus simulans* | 3,10E+07 | 7,10E+06 | 1,09E+04 |
| F | 12_05_23 | *Pseudomonas aeruginosa* | 3,79E+07 | 1,39E+07 | 3,54E+04 |
| F | 12_05_23 | *Pseudomonas aeruginosa* | 4,22E+07 | 7,50E+06 | 3,42E+04 |
| F | 12_05_23 | *Pseudomonas aeruginosa* | 3,29E+07 | 1,71E+07 | 1,74E+04 |
| F | 12_05_23 | *Proteus mirabilis* | 1,64E+07 | 1,97E+07 | 2,77E+04 |
| F | 12_05_23 | *Proteus mirabilis* | 2,79E+07 | 1,97E+07 | 3,59E+04 |
| F | 12_05_23 | *Proteus mirabilis* | 3,75E+07 | 3,07E+07 | 3,75E+04 |
